# Integrative processing in artificial and biological vision predicts the perceived beauty of natural images

**DOI:** 10.1101/2023.05.05.539579

**Authors:** Sanjeev Nara, Daniel Kaiser

## Abstract

Previous research indicates that the beauty of natural images is already determined during perceptual analysis. However, it is still largely unclear which perceptual computations give rise to the perception of beauty. Theories of processing fluency suggest that the ease of processing for an image determines its perceived beauty. Here, we tested whether perceived beauty is related to the amount of spatial integration across an image, a perceptual computation that reduces processing demands by aggregating image elements into more efficient representations of the whole. We hypothesized that higher degrees of integration reduce processing demands in the visual system and thereby predispose the perception of beauty. We quantified integrative processing in an artificial deep neural network model of vision: We compared activations between parts of the image and the whole image, where the degree of integration was determined by the amount of deviation between activations for the whole image and its constituent parts. This quantification of integration predicted the beauty ratings for natural images across four studies, which featured different stimuli and task demands. In a complementary fMRI study, we show that integrative processing in human visual cortex predicts perceived beauty in a similar way as in artificial neural networks. Together, our results establish integration as a computational principle that facilitates perceptual analysis and thereby mediates the perception of beauty.

## Introduction

During our daily lives, some visual images reliably evoke a feeling of beauty while others do not. But why do images differ in their ability to evoke beauty? Recent research suggests that perceptual factors play a critical role. Studies on low-level visual features revealed a set of properties that are associated with perceived beauty, such as an image’s color, curvature, or symmetry (Enquist and Arak, 1994; Palmer et al., 2013; Schloss and Palmer, 2011; Silvia and Barona, 2009). The perception of beauty may thus arise from the presence of relatively basic features, as well as their spatial configuration in the image (Palmer et al., 2013; Van Geert and Wagemans, 2020). The important role of visual image properties in evoking beauty is consistent with neuroscientific studies that show that beauty ratings for natural images are predicted by activations in cortical regions responsible for perceptual analysis (Vessel et al., 2019; Yue et al., 2007; Zhao et al., 2020) and early neural responses associated with perceptual processing (Kaiser, 2022a).

Despite the realization that the beauty of an image is – at least partially – determined during perceptual analysis, it is still largely unclear which perceptual mechanisms govern the formation of the phenomenological experience of beauty. An intriguing proposal is that processing fluency, that is the ease with which is stimulus can be analyzed in the visual system, plays a critical role for perceived beauty (Forster, 2020; Reber et al., 2004). Critical evidence for this proposal comes from studies that have shown that the goodness of a composition (such as whether multiple image element from a meaningful whole) impacts both perceptual processing efficiency and perceived beauty (Van Geert and Wagemans, 2020; Palmer et al., 2013).

On the neural level, typical compositions were previously associated with a release from neural competition, rendering it easier to efficiently represent multiple stimuli at the same time (Kaiser et al., 2014a; McMains and Kastner, 2010). This decrease in competition has been linked to an increasing capacity for integration: When image elements (such as multiple objects) can be represented as a meaningful whole, rather than multiple unrelated entities, the visual system can efficiently reduce the complexity of representations (Kaiser et al., 2019, 2014a). Although we know that efficient neural integration facilitates perceptual analysis, it is unclear whether efficient integration similarly determines perceived beauty.

Here, we thus tested whether the degree of visual integration across an image can reliably predict whether natural images are perceived as beautiful. To quantify integration, we used a deep neural network (DNN) as a model of the biological visual processing cascade (Cichy and Kaiser, 2019; Kriegeskorte, 2015). Within this neural network, we quantified integration by computing how well whole images were predicted by a combination of their individual parts. This integration measure predicted perceived beauty across four studies with different natural images and under different task demands. Applying the same analysis logic to human fMRI data recorded for a subset of images, we show that integration in scene-selective visual cortex predicts perceived beauty in a similar way as our DNN-derived quantification of integration. Together, these results highlight that integration is a critical computation for the evaluation of beauty in natural images.

## Results

Here, we tested whether the degree of visual integration across an image can reliably predict whether the image is rated as beautiful. To quantify integration, we used a deep neural network (DNN) as a model of the biological visual system (Cichy and Kaiser, 2019; Kriegeskorte, 2015). Specifically, we used a VGG16 network architecture (Simonyan and Zisserman, 2015) pretrained on scene categorization using the Places365 image set (Fig. 1a; Zhou et al., 2017).

**Figure 1.**
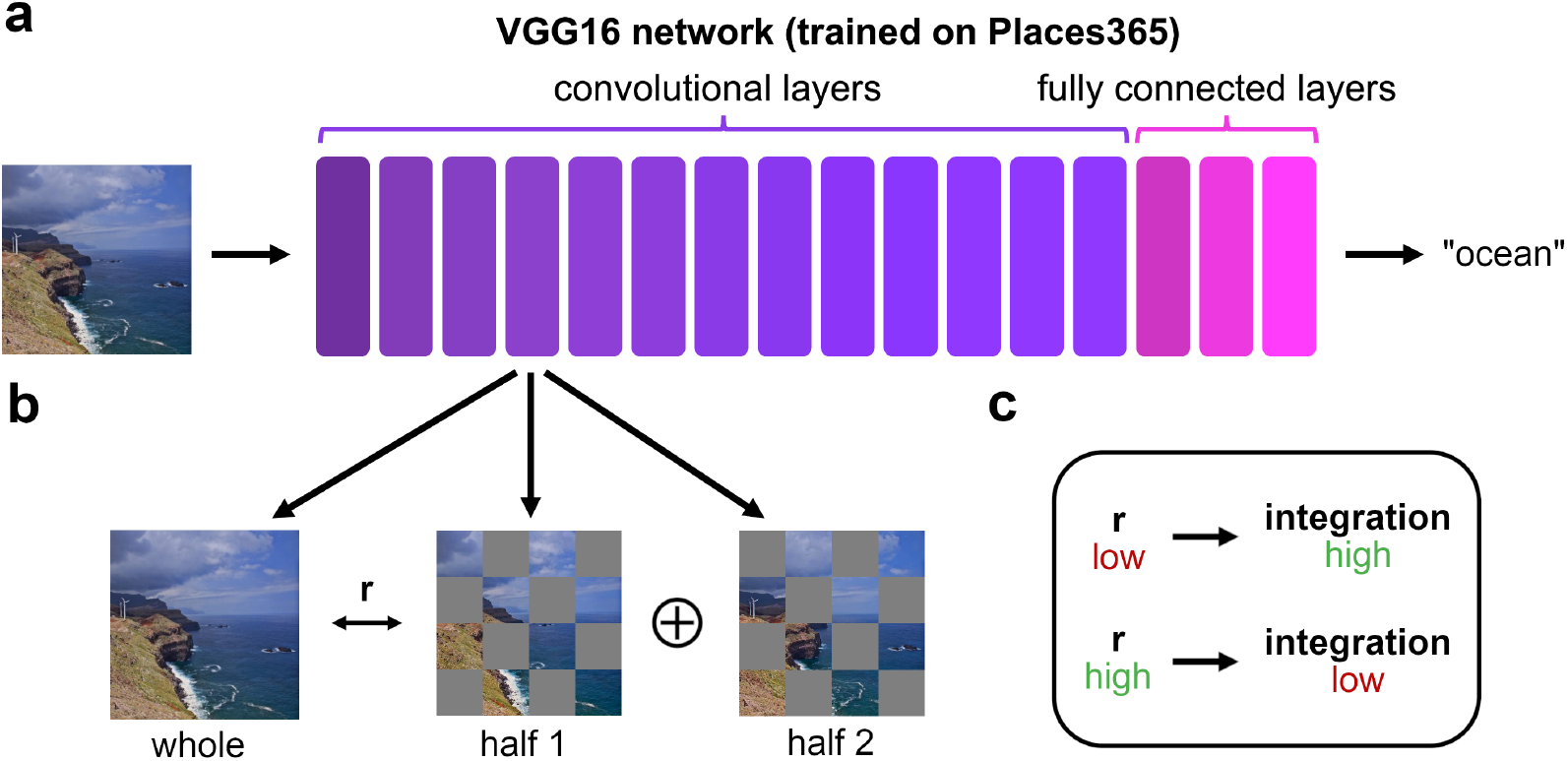
DNN-based quantification of integrative processing. **a)** We used a VGG16 DNN as an in-silico model of cortical scene processing. The DNN was trained on scene categorization using the Places365 dataset. **b)** We fed the DNN with each full image, as well as with two halves of the images. Halves were generated by obscuring 50% of the image in a checkerboard-like fashion, with different spatial scales (i.e., number of squares on the checkerboard). The example shows the 4×4 spatial scale. To quantify integration, we correlated the layer-specific activation pattern to each whole image with the average of the activation patterns to the two halves. **c)** When the resulting correlation is low, one can infer more integration (as integrative processes are not captured by the activation patterns to the parts), whereas when the correlation is high, one can infer less integration (as the average of the parts accurately captures the whole).

Within this neural network, we quantified integration by computing how well whole images were predicted by a combination of their individual parts, where accurate prediction indexes largely parallel processing, and thus a low degree of integration, while less accurate prediction indexes interactive processing across image parts, and thus a higher degree of integration. This logic is derived from human neuroimaging studies, which have shown that the average response to multiple image elements accurately predicts the response to the full image (Jeong and Xu, 2017; Kaiser et al., 2014b; Kaiser and Peelen, 2018; Kliger and Yovel, 2020; MacEvoy and Epstein, 2009), and that a relative decrease in this prediction indicates integrative processing that renders responses to the whole image dissimilar from the responses to its parts (Kaiser et al., 2019; Kaiser and Peelen, 2018; Kubilius et al., 2015). Recent computational investigations suggest that DNNs show a similar averaging of responses for multiple image elements (Jacob et al., 2021), although average responses in DNNs do not always resemble representations to the whole as faithfully as in the human brain (Mocz et al., 2023).

Here, we computed integrative processing for individual images, by feeding the network two halves of an image (e.g., the bottom-left and top-right quadrants versus the bottom-right and top-left quadrants), as well as the full image (Fig. 1b). For each image, and separately for each network layer, we then computed how much the activation pattern to the full image was correlated to the average activation pattern to the two halves. The strength of this correlation was taken as a measure of integrative processing, where lower values (i.e., a higher dissimilarity between the whole and its parts) indicates a higher degree of integration (Fig. 1c). The integration measure was computed separately at 5 different spatial granularities, where halves were created by dividing the image into 2×2, 4×4 (as in Fig. 1b), 8×8, 16×16 or 32×32 identical squares. Each image half contained all odd or all even squares (i.e., corresponding to either all white or black squares on a checkerboard). This procedure allowed us to probe integrative processing at different spatial scales.

We then tested whether our DNN-derived integration measure could successfully predict perceived beauty. To this end, we use the image-specific integration measure to predict beauty ratings in a series of four studies with varying image sets and task demands.

In Study 1, we collected beauty ratings for 250 natural scene images from 25 online participants (Fig. 2a; see Materials and Methods for details). During our experiment, we only showed the images briefly (50ms exposure time). Previous work has shown that observers can judge the beauty of an image even under such brief presentation regimes (Verhavert et al., 2018). We reasoned that brief exposure would lead to more successful prediction of perceived beauty from our integration measure, as observers do not have time for extensive cognitive evaluation of the image. Correlating the integration measure with beauty ratings revealed a strong relationship between integration and perceived beauty, with correlations of up to r=0.6 (Fig. 2b), showing that a higher degree of integrative processing predisposes a higher beauty rating. There were two notable patterns in these correlations: First, correlations were strongest in intermediate to late network layers, suggesting that integration over mid- and high-level features determines perceived beauty. Second, correlations were apparent across all spatial scales, but strongest for the coarser scales, with a decrease for the 16×16 and 32×32 scales. This suggests that integration across larger parts of the images is a stronger predictor of perceived beauty than integration across fine details. To test whether the integration measure could also accurately predict beauty ratings for novel images, we next fitted a linear model on integration values derived from all 16 network layers, using all but one image. We then predicted the values for the left-out image using the fitted model. After each image was left out once, we correlated the model predictions with the real beauty ratings. We found that the linear model could successfully predict beauty ratings for novel images at all spatial scales (Fig. 2c), with numerically strongest predictions for an intermediate 8×8 spatial scale.

**Figure 2.**
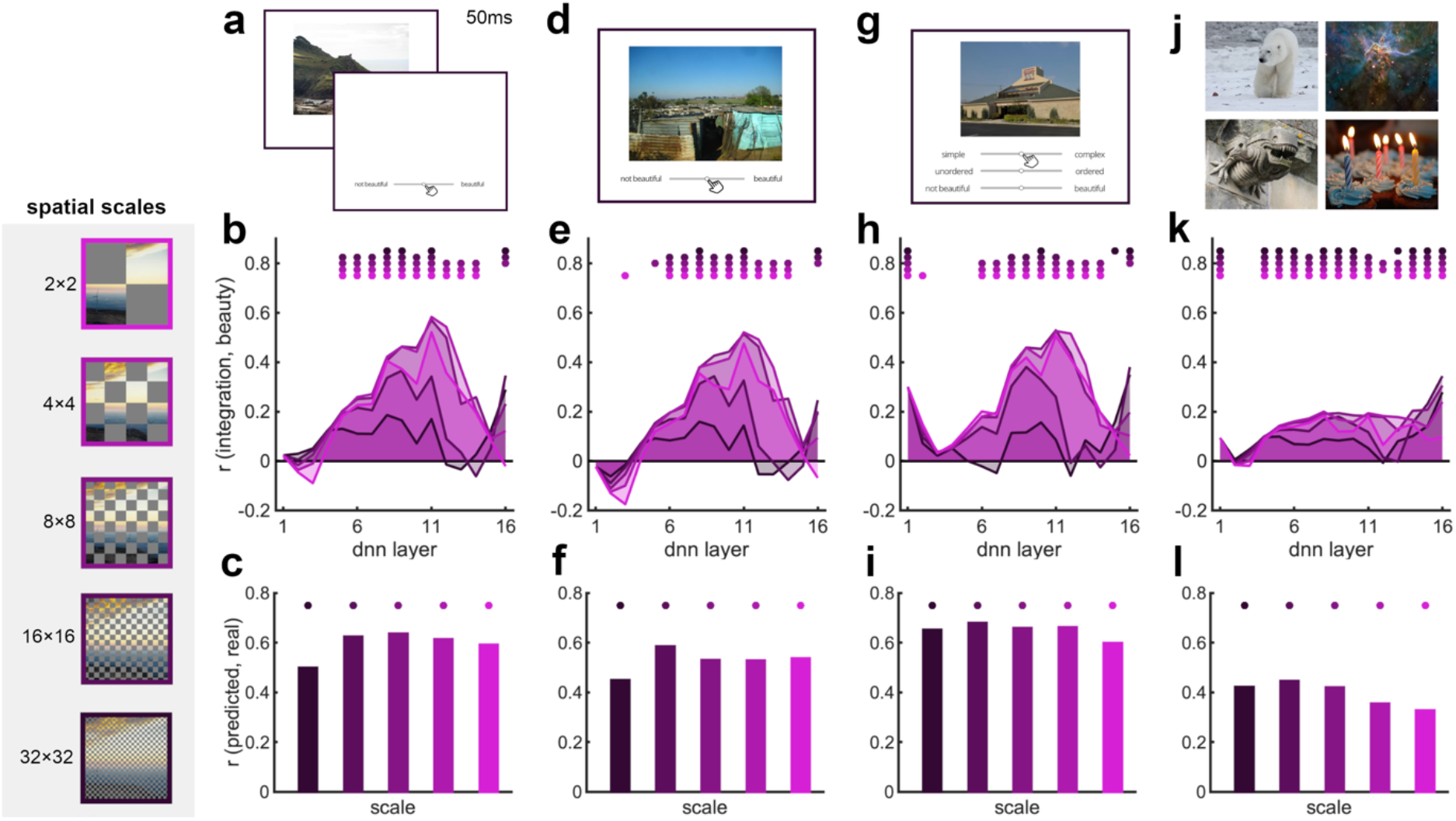
The degree of integrative processing in a DNN predicts perceived beauty. **a)** In Study 1, participants briefly viewed 250 natural scene images and rated their beauty. **b)** We then correlated the DNN-derived integration measure (see Fig. 1) with beauty ratings across images, separately for each network layer and each spatial scale (see left). Integration significantly predicted beauty ratings across images, with highest correlations in intermediate network layers and coarser spatial scales. **c)** To assess predictions for novel images, we further estimated linear models with the beauty ratings as the criterion and the integration measure in each layer as predictors, for all but one image. We then generated predictions for the left-out images with these linear models and correlated the predicted ratings of the left-out images with the real ratings for these images. The linear model could predict ratings for novel images across all spatial scales. **d)** In Study 2, participants viewed the same set of images as in Study 1, now with unlimited viewing time. **e/f)** Results strongly resembled the results from Study 1. **g)** In Study 3, participants viewed a disjoint set of 250 natural scenes images, again with unlimited viewing time. Here, participants additionally rated the images’ complexity and order (see text). **h/i)** Results were again similar to Studies 1 and 2. **j)** In Study 4, we used beauty ratings collected for 900 vastly different natural images from the OASIS database. **k/l)** Even for such diverse images, our integration measures significantly predicted beauty ratings, though to a lesser extent than for the more homogeneous natural scene images. Dots indicate p<0.05 (corrected for multiple comparisons).

In Study 2, we asked whether the same pattern of results could be replicated when observers are able to study the image as long as they want. We thus tested another 25 online participants, who rated the same 250 natural scene images. Here, they had unlimited time to provide their beauty rating while the image stayed on the screen, allowing them to also cognitively evaluate the image in detail (Fig. 2d). Perhaps surprisingly, the integration measure essentially predicted perceived beauty equally well as for briefly presented images (Fig. 2e/f), replicating the pattern obtained in Study 1.

In Study 3, we sought to replicate the findings from Study 2 with a completely new set of 250 natural scenes, rated for their beauty by 26 online observers. Here, observers additionally rated the complexity and order of each image (Fig. 2g; see Materials and Methods for details), which allowed us to gauge how the integration measure relates to human ratings of how complex or ordered an image is (see below). Results again replicated the pattern from Studies 1 and 2, showing that integrative processing is a strong predictor for perceived beauty (Fig. 2h/i). We further tested whether human-rated image complexity and order could explain integration within the DNN. Although complexity and order both linearly predicted beauty ratings (r=0.11, p=0.09 for complexity, r=0.31, p<0.001 for order), they were only moderately correlated to the integration measure derived from the DNN (all r<0.22 for complexity and r<0.22 for order). When complexity and order ratings were partialed out, integration in the DNN could still predict beauty ratings well (see Supplementary Fig. S1), suggesting that human ratings of complexity and order do not fully capture the visual features that predispose integrative processing in the DNN.

Finally, in Study 4, we tested whether our integrative processing measure could predict perceived beauty not only for natural scenes, but for a wide range of photographs that depict objects, people, and everyday situations. Here, we utilized beauty ratings obtained for a set of 900 diverse natural images contained in the OASIS database (Fig. 2j; (Brielmann and Pelli, 2019; Kurdi et al., 2017). For these images, given their large variability in content and emotional valence, we expected that predictions derived from our integrative processing measure would be reduced. If the integration measure still predicted perceived beauty in these images, however, it suggests that integrative processing is a computation that predisposes beauty across natural images from various domains. As hypothesized, correlations were indeed lower, but integrative processing still predicted beauty ratings (Fig. 2k). Similar to the previous studies, perceived beauty was again better predicted from intermediate layer activations and from coarser spatial scales. Despite these reduced correlations, the DNN-derived integration measure still successfully predicted beauty ratings for novel images (Fig. 2l).

In supplementary analyses, we show that the same prediction of perceived beauty can be achieved with a VGG16 network trained on object categorization instead of scene categorization (Supplementary Fig. S2). We further evaluate a second possible predictor of perceived beauty: We evaluated part-based similarity, that is the degree to which parts of an image show visual similarity to other parts of the image. Previous research has suggested that similarity between multiple parts can – like integration across parts – alleviate neural competition in visual cortex (Beck and Kastner, 2007). To quantify part-based similarity, we computed the similarity in DNN activations to one half of the image and the other half of the image. Our data show that part-based similarity is also capable of predicting perceived beauty, albeit to a lesser extent than integration (Supplementary Fig. S3a/b) and that the association between integration and beauty cannot be explained by an image’s self-similarity (Supplementary Fig. S3c). Finally, we show that the degree to which whole images activate the network (operationalized through the L2 norm of the activation pattern in each layer) also predicts beauty ratings to some extent, but that this prediction cannot account for the – stronger – predictions provided by our integration measure (Supplementary Fig. S4).

Together, our four studies demonstrate that a DNN-derived measure of integrative processing predicts perceived beauty across different natural images and under different task demands. Using a DNN provides a powerful way of estimating integrative processing in an objective and scalable way, which can be used to derive a computational prediction for perceived beauty across large sets of images.

However, we cannot be fully sure whether our DNN indeed replicates the way in which the human brain integrates information across images: After all, DNNs have been shown to diverge from human visual processing in potentially critical ways (Bowers et al., 2022; Mocz et al., 2023; Xu and Vaziri-Pashkam, 2021). Given this backdrop, we wanted to investigate whether an integrative processing measure derived from human fMRI data is similarly capable of predicting perceived beauty. We thus ran an fMRI study, in which 21 participants viewed a set of 32 natural scene images (which were rated for beauty in Study 3). During this study, participants viewed the whole images as well as their two complimentary halves (see Materials and Methods for details). Halves were the bottom-left and top-right versus the bottom-right and top-left quadrants, thus resembling the 2×2 spatial scale in the DNN analysis. We extracted multi-voxel fMRI patterns from a set of regions in retinotopic early visual cortex (V1, V2, V3, and V4) and scene-selective cortex (occipital place area - OPA, medial place area - MPA, and parahippocampal place area - PPA). We then performed an analysis analogous to our DNN analysis: For each scene, we computed the correlation between the multi-voxel response pattern to the whole image and the average of the multi-voxel response pattern to the two halves (Fig. 3a; see Materials and Methods for details). This correlation was again used as a measure of integrative processing, where lower correlations indicate a higher degree of integration (Kaiser et al., 2019; Kaiser and Peelen, 2018; Kubilius et al., 2016).

**Figure 3.**
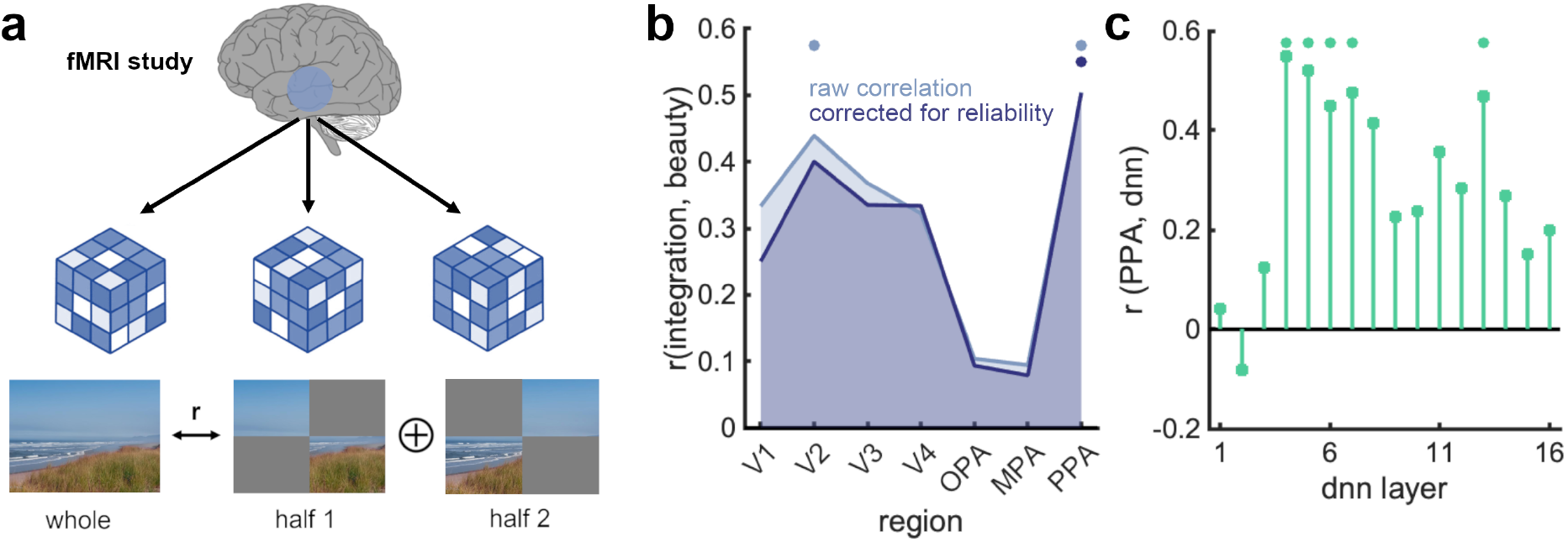
Integrative processing in scene-selective PPA predicts perceived beauty. **a)** To test whether integrative processing in the human visual system similarity predicts perceived beauty as integrative processing in an artificial DNN, we conducted an fMRI study in which participants viewed 32 whole natural scene images and their corresponding halves (on the 2×2 spatial scale). Integration was quantified by computing the correlation between the response pattern evoked by each whole image (from half of the runs) and the average of the response patterns evoked by its two halves (from the other half of runs). Integration was assessed for four regions in early visual cortex (V1-V4) and four regions in scene-selective cortex (OPA, MPA, and PPA). **b)** The degree of integrative processing in V2 and PPA significantly correlated with beauty ratings across images, suggesting that integrative processing in the biological visual system is similarity related to perceived beauty as integrative processing in an artificial DNN. The correlation between integration and beauty ratings in PPA remained significant when controlling for the reliability of fMRI responses to the whole images (by assessing correlations in response patterns evoked by the whole images across experimental runs). **c)** Across images, the degree of integrative processing in the DNN (on the 2×2 spatial scale) correlated with the degree of integration in the PPA, specifically in middle layers of the DNN, revealing a correspondence between integration in biological and artificial vision. Dots indicate p<0.05 (corrected for multiple comparisons).

Correlating the integration measure derived from the fMRI data with the beauty ratings for the 32 images used in the fMRI experiment revealed a significant correlation in V2 and scene-selective PPA (Fig. 3b). However, there is a possibility that different degrees of image-specific integration simply reflect differences in the reliability of responses across images: If an image yields less reliable responses, the response cannot be approximated well to begin with. To address this concern, we additionally computed a measure of reliability for each whole scene (see Materials and Methods), which we partialed out when correlating the integration measure and the beauty ratings. In this analysis, integration in PPA still significantly predicted perceived beauty. We also correlated the integration measure obtained from the PPA with the integration measure obtained from the DNN on 2×2 spatial scale (Fig. 3c). We found significant correlations between the degree of integrative processing in the PPA and the DNN, again peaking at the intermediate layers, where integration was most predictive of the beauty ratings (see Fig. 2). This suggests that integration varies across images in similar ways in human visual cortex and in our DNN model.

Together, our fMRI results show that integrative processing in scene-selective cortex, and specifically in the PPA, predicts the aesthetic appeal of natural images. Together with the DNN analyses, our fMRI data thus provide converging evidence that spatial integration is a critical computation that predisposes perceived beauty.

## Discussion

Our study unveils that integrative processing, as a computational principle in the visual system, is predictive of the perceived beauty of natural images. We show that a quantification of integration in an artificial neural network can reliably predict beauty ratings for natural scene images across different scene image sets and across different stimulus presentation times. Even when extrapolating the analysis to a widely varying set of natural images (containing objects, people, and scenes), integration in the network could predict perceived beauty. In an fMRI analysis, we show that an analogous quantification of integration in the human visual system can equally predict beauty ratings for natural scenes. Together, these results provide critical evidence for the notion that efficient processing in sensory systems, as enabled by efficient spatial integration, is linked to the aesthetic evaluation of sensory inputs.

This notion fits well with theories that explain perceived beauty through processing fluency in brain systems (Forster, 2020; Oppenheimer, 2008; Reber et al., 2004). Efficient integration has been previously related to a decrease in neural competition between image elements, both in simple (Kubilius et al., 2015; McMains and Kastner, 2010) and naturalistic (Kaiser et al., 2014a; Kaiser and Peelen, 2018) visual stimuli. On this note, integration reduces interference between image elements by aggregating them into fewer compound representations (Kaiser et al., 2019, 2014a) and thereby increases the ease with which these fewer representations can be processed in parallel. Integrative processing may of course not constitute the only computational principle that determines processing fluency, and thereby perceived beauty. Uncovering other complementary principles may further increase the amount of variance in beauty ratings that can be predicted from sensory-derived measures of stimulus processing.

Our DNN results further highlight that integration effects at different spatial scales can predict perceived beauty. This suggests that the efficiency of sensory processing that is critical for rating aesthetic appeal is computed at different levels of detail, from the integration of local spatial features that do not necessarily transport semantic information to the integration of larger regions that unites semantically meaningful image elements. It will be interesting to see whether integration effects at different scales relate to different processing steps in the visual hierarchy, with varying sensitivity to visual and semantic information. An interesting future avenue would be to not only compute the integration of purely visual information: For instance, future studies could also look at conceptual descriptors (e.g., via using language models; Bonner and Epstein, 2021; Hayes and Henderson, 2021) to capture information integration in conceptual space for visual images.

Our fMRI results show that integrative processing in the biological brain can predict perceived beauty in a similar way as integrative processing in DNNs. This finding further stresses the mechanistic similarity between DNNs and the human visual system (Kriegeskorte, 2015; Kietzmann et al., 2019; Lindsay, 2021) and showcases the possibility to use DNNs as high-throughput tools for uncovering the computational principles that guide visual processing (Cichy and Kaiser, 2019; Wichmann and Geirhos, 2023). Importantly, the focus on principles of processing rather than on pure predictive accuracy (which has been demonstrated elsewhere; Conwell et al., 2021; Jin et al., 2019; Lu et al., 2014; Seresinhe et al., 2017) opens the door to uncovering – and subsequently validating – computational implementations in the biological brain. This computational approach promises to reveal other important computational principles that govern the sensory stages of aesthetic appreciation.

In the human visual system, the beauty of natural scene images was only robustly predicted from responses in scene-selective PPA. Previous studies indicate that PPA is capable of integrating scene elements distributed across visual space, specifically when the scenes form a meaningful perceptual whole (Chen et al., 2023; Kaiser et al., 2020; Kaiser and Cichy, 2021). Here, we show that such integration processes systematically vary across scenes, and that this variation is a determinant for a scene’s aesthetic appeal. An open question concerns what exactly is integrated in PPA. Processing in this region has previously been linked to both categorical and semantic attributes of a scene (Epstein and Baker, 2019; Harel et al., 2013; Walther et al., 2009); as well as to low- and mid-level visual properties that are reliably associated with scenes (Nasr et al., 2014; Watson et al., 2017, 2016). Our DNN analysis suggest that both mid- and high-level properties computed at intermediate to deep layers may be critical for determining perceived beauty, and PPA activations have indeed been linked to activations in intermediate to deep DNN layers before (Dwivedi et al., 2021; Groen et al., 2018). At this point, more research is needed to uncover the critical features across which the PPA integrates information. One notable limitation of the current fMRI study is that we could only evaluate a small subset of images. Besides the PPA, our data show substantial correlations between integration and beauty ratings, which did not reach statistical significance given the limited number of stimuli. It would thus be premature to dismiss a link between integrative processing in early visual cortex and perceived beauty.

Finally, our study quantified integration across visual space. Yet, space is not the only dimension across which efficient integration can mediate perceived beauty. For example, a recent study used DNN models to quantify the integration of visual features across hierarchical levels to model aesthetic perception, and has in turn linked such hierarchical integration processes to parietal and frontal brain systems (Iigaya et al., 2023). Future studies could also link perceived beauty to integration across time: Recent studies in neuroaesthetics increasingly focus on more naturalistic and dynamic stimuli (Isik and Vessel, 2021; Kaiser, 2022b; van Elk et al., 2019; Zhao et al., 2020), and it will be interesting to see whether efficient information integration in the time domain can also predict the aesthetic appeal of dynamically evolving visual scenes.

In sum, our study establishes integrative processing as a computational principle in the visual system that is capable of explaining perceived beauty. When images are more strongly integrated across visual space – and consequentially the representation of the whole gets more dissimilar to the representation of its parts – they are perceived as more beautiful. Gestalt psychologists famously noted that “the whole is something else than the sum of its parts” (Koffka, 1935). Here we show that the degree to which the whole is different from its parts not only has an impact on the formation of efficient representations the whole but that this process is linked to whether or not we assign beauty to it.

Our discovery provides a new impulse for research in neuroaesthetics: Moving from the study of individual visual features and their impact on aesthetic appeal towards the study of overarching computational principles has the potential to reconcile research on different visual features and stimulus domains.

## Materials and Methods

### Participants

We ran three online studies in which participants rated the beauty of natural scene images. Study 1 was completed by 25 participants (mean age 23.9, SD=5.2; 18 male, 7 female), Study 2 was completed by 25 participants (mean age 24.0, SD=4.3; 10 male, 15 female), and Study 3 was completed by 26 participants (mean age 24.9, SD=5.2; 12 male, 13 female, 1 non-binary). Participants were recruited through Prolific (www.prolific.co) and received monetary reimbursement. Informed consent was provided through an online form. The studies were approved by the Ethical Committee of the Institute of Psychology at the University of York.

We additionally ran an fMRI study, where participants viewed whole and partial natural images. The fMRI study was completed by 22 participants (mean age 28.8, SD=4.5; 10 male, 12 female). One participant did not complete all experimental runs and was excluded from further analyses. Participants received monetary reimbursement and provided written informed consent. The study was approved by the General Ethical Committee of the Justus-Liebig-University Gießen.

Sample sizes were determined through convenience sampling, as data were subsequently averaged across participants for our analyses. Reliability measures showed good agreement between participants (see below).

### Rating study design

Three behavioral studies were conducted online, using the Gorilla platform for online testing (Anwyl-Irvine et al., 2020).

In Study 1, participants viewed 250 images of natural scene photographs in random order, sampled from the validation set of the Places365 dataset (Zhou et al., 2017) to include a large variety of contents. All images depicted outdoor scenes. On every trial, they viewed a single scene, presented for 50ms. After the scene presentation, they were presented with a slider, operated by the mouse, on which they adjusted how beautiful they rated the scene. Slider values were coded between 0 and 100. Before the experiment, beauty was defined to participants as how beautiful or aesthetically pleasing the scene is. After participants provided their ratings, participants could advance to the next trial by pressing a mouse button.

In Study 2, participants viewed the same 250 images as in Study 1, in random order. The study design was identical to Study 1, but here the slides was shown at the same time as the scene (below the image). The scene was visible until participants provided their rating. Participants were not instructed to respond fast and were given as much time as needed to adjust the slider.

In Study 3, participants viewed another set of 250 images in random order, sampled similarly to Studies 1/2, but the image set was completely disjunct from the previous set. The design was identical to Study 2, but on every trial the scene was accompanied by three sliders, on which they could adjust: (1) how complex they rated the scene, (2) how ordered the rated the scene, and (3) how beautiful they rated the scene. Complexity was defined as the number of distinct elements (such as objects, shapes, or colors) present in the scene - compared to how many such items are expected in a typical scene. Order was defined as the degree to which the different scene elements (such as objects) are positioned across the scene and relative to each other in a typically structured manner. Both complexity and order predicted beauty ratings linearly (see results).

### Database ratings

In addition to the three behavioral studies, we used beauty ratings collected for the images in the OASIS database (Kurdi et al., 2017). This database contains 900 photographs depicting a large variety of contents, from people to objects and scenes. Beauty ratings for these images from a total of 757 observers were compiled by Brielmann and Pelli (Brielmann and Pelli, 2019). In their study, each individual image was rated by at least 104 observers.

### Behavioral data analysis

For Studies 1-3, as well as the OASIS beauty ratings, a mean score for each image was computed from the ratings of all observers. Beauty ratings were reliable across people in Studies 1-3, as shown by split-half correlations (r>0.89 for all studies). Detailed reliability measures for the OASIS ratings are reported elsewhere (with split-half reliability r>0.95; Brielmann and Pelli, 2019).

### Deep neural network analysis

We used a VGG16 (Simonyan and Zisserman, 2015) deep neural network (DNN) trained on scene categorization using the Places365 image set (Zhou et al., 2017). The VGG architecture was chosen as it has been shown to provide a good computational approximation of the ventral visual pathway (Schrimpf et al., 2021). The pre-trained DNN was obtained from https://github.com/CSAILVision/places365. The network was originally deployed in Caffe (Jia et al., 2014) and imported to Matlab using the Caffe Importer for the Matlab Deep Learning Toolbox (https://de.mathworks.com/matlabcentral/fileexchange/61735-deep-learning-toolbox-importer-for-caffe-models). Results from a VGG16 network trained on Imagenet (Deng et al., 2009; implemented in Matlab) are reported in Supplementary Fig. S2.

We fed the network with either the full scene or two halves of the scene. Halves were created by slicing the scene into 2×2, 4×4, 8×8, 16×16 or 32×32 identical squares. Each image half contained all odd or all even squares (i.e., corresponding to either all white or black squares on a checkerboard). This slicing yielded two image halves for each of 5 different spatial scales. The full image and all possible halves were fed to the network, and layer-specific activation patterns for each image were obtained separately for all 16 layers of the network.

To quantify integration, we correlated the layer-specific activation patterns to the whole scene with the average response pattern to the scene halves, separately for each spatial scale (e.g., by slicing into 2×2 pieces). Note that any linear combination with equal weights (e.g., the sum) would yield equivalent correlations.

The sign of the resulting correlations was flipped to yield a measure of integration: When processing is largely parallel, activation patterns to the full image should be accurately predicted by the average of the activation patterns to the two halves (resulting in a low integration measure), as previously shown in human cortex (Jeong and Xu, 2017; Kaiser et al., 2014b; Kaiser and Peelen, 2018; Kliger and Yovel, 2020; MacEvoy and Epstein, 2009) and for DNNs (Jacob et al., 2021; Mocz et al., 2023). Unlike in human cortex, averaging the response patterns to constituent objects does not perfectly predict the response to multiple objects (Mocz et al., 2023) – in our analysis, however, the overall quality of the fit is not critical, as we only examine relative differences in the fit across images. When processing is more integrative, activation patterns to the full image should be less accurately predicted by the average of the activation patterns to the two halves (resulting in a higher integration measure). Measuring integration through such multivariate pattern combination analysis has successfully been employed in fMRI work (Baldassano et al., 2017; Kaiser et al., 2019; Kaiser and Peelen, 2018; Kubilius et al., 2015). In sum, our procedure thus yielded a quantification of integration for each image, at each spatial scale, and in each network layer.

We additionally computed a part-based similarity measure as an alternative predictor for perceived beauty. To quantify part-based similarity, we correlated the layer-specific activation patterns to the two halves, separately for each spatial scale. This correlation can be directly interpreted as a measure of part-based similarity, where higher correlations signal greater visual correspondence between the image halves. Results for the part-based similarity measure are reported in Supplementary Fig. S3.

Finally, we computed how much each whole image drives the DNN, as another alternative predictor for perceived beauty. Here, we computed the L2 norm for each network layer as a measure of activation strength. The relationship of the L2 norms and the beauty ratings are reported in Supplementary Fig. S4.

### Predicting beauty ratings from DNN integration

To assess how beauty rating were predicted by integration in the DNN, we correlated (Spearman-correlations) the image-specific mean beauty ratings for each of the four studies with the image-specific quantifications of integration, separately for each network layer. The same analysis was performed for the part-based similarity measure, as reported in Supplementary Fig. S3. All p-values corresponding to these correlations were false-discovery-rate corrected across network layers and spatial scales.

To assess the unique contribution of integration in predicting beauty ratings, we also performed partial correlation analyses, where either the part-based similarity measure or the activation strength for the whole image was partialled out when correlating the beauty ratings with the integration measure. These analyses are reported in Supplementary Fig. S3/S4.

We also assessed whether the DNN integration measure could successfully predict beauty ratings for novel images. To this end, we fitted a linear model with all layers included as predictors for the beauty ratings for all images but one. Both the criterion and predictors were z-scored prior to estimating the regression weights. We then derived a predicted beauty rating for the left-out image from the estimated linear model.

Repeating this procedure for all images being left out once yielded a predicted value for each image. To assess how well the model could predict the beauty ratings for the held-out images, we correlated the predicted beauty ratings with the real beauty ratings obtained from our human observers.

### fMRI study

During the fMRI experiment, participants viewed 32 scene images, which were a subset of the stimulus set used in Study 3. These scenes were chosen to come from four broadly defined categories (beaches, buildings, highways and mountains) and to cover a range of beauty ratings (mean average rating: 66/100; minimum: 35; maximum: 87). The scenes were cropped so that all images had square aspect ratio and resized to 512×512 pixels. In each of ten fMRI runs (4.5min each), participants completed 128 trials (1280 in total). In each trial, they saw a scene image (7.5×7.5 degrees visual angle) for 500ms, followed by an inter-trial interval of 1500ms, during which a black fixation cross was shown. During each run, each scene was shown in three possible conditions: whole image (32 trials per run), top-left and bottom-right quadrants only (32 trials per run), or top-right and bottom-left quadrants only (32 trials per run). These conditions thus corresponded to the 2×2 spatial scale in the DNN analysis. Each run additionally featured 32 fixation trials, during which the black fixation cross turned gray. Participants were instructed to press a button on these trials.

### fMRI acquisition and preprocessing

MRI data was acquired using an 3T Siemens Magnetom PRISMA Scanner equipped with a 64-channel head coil. T2*-weighted gradient-echo echo-planar images were collected as functional volumes (TR=1850ms, TE= 30ms, 75° flip angle, 2.2mm^3^ voxel size, 58 slices, 20% gap / distance factor, 220mm FOV, 100×100 matrix size, interleaved acquisition). Additionally, a T1-weighted image (MPRAGE; 1mm^3^ voxel size) was obtained as a high-resolution anatomical reference. During preprocessing, the functional volumes were realigned and coregistered to the T1 image using SPM12 (www.fil.ion.ucl.ac.uk/spm/). The functional data were then modelled using a general linear model with separate predictors for the 32 images and the three presentation conditions (the whole image and the two parts), separately for each run. The GLM also contained six movement regressors obtained during realignment, and thus 102 regressors for each run.

### fMRI analyses

Multivariate analyses were performed using the CoSMoMVPA toolbox (Oosterhof et al., 2016). Multi-voxel response patterns were obtained for seven regions of interest. Four early visual cortex regions were defined using a template atlas (Wang et al., 2015): V1, V2, V3, and V4. Additionally, three scene-selective regions were defined using functional group maps (Julian et al., 2012): the occipital place area (OPA; also termed transverse occipital sulcus), the medial place area (MPA; also termed retrosplenial cortex), and the parahippocampal place area (PPA). For each region, multi-voxel response patterns were extracted by unfolding the GLM beta weights for each image and each run into a one-dimensional vector.

We then performed an analysis similar to the DNN analysis. For each region, we averaged response patterns evoked by the two halves and correlated the resulting response pattern to the response pattern evoked by the full image, separately for each image. Here, the response pattern to the whole scene always stemmed from half of the fMRI runs and the average response pattern to the halves stemmed from the other half of runs (Kaiser and Peelen, 2018). Correlations were computed across all possible 50/50 splits among the 10 runs and averaged across splits. By flipping the sign of the resulting correlations, we obtained a quantification of integration, where lower correlations index a higher degree of integration. These values were averaged across participants before comparing them to the beauty ratings.

From the fMRI data, we additionally obtained a quantification of response reliability for the whole images. This was done by correlating the response pattern to each whole image in half of the runs with the response pattern to the same image in the other half of runs. Correlations were again averaged across all possible 50/50 splits among the 10 runs. This yielded a correlation for each whole image that indexed the stability of the response patterns across repetitions.

### Predicting beauty ratings from fMRI integration

To assess how beauty rating were predicted by integration in the human visual cortex, we correlated (Spearman-correlations) the image-specific mean beauty ratings for the 32 images used in the fMRI study with the image-specific fMRI quantifications of integration, separately for each brain region. All p-values corresponding to these correlations were false-discovery-rate corrected across regions.

We additionally repeated this analysis while partialing out the quantification of reliability obtained for each image. In this way, we could ensure that differences in integration (i.e., differences in how well the combined response to the image halves predicted the response to the whole image) were not simply resulting from differences in the reliability of responses to each of the individual images (where less reliable response would be harder to approximate in the first place).

## Data availability

Data, code, and materials are available on the Open Science Framework (OSF), at https://osf.io/87zxp/. Raw and preprocessed fMRI data are available on request, as they are also analyzed for another project.

## Acknowledgements

D.K. is supported by the Deutsche Forschungsgemeinschaft (DFG; SFB/TRR135 – “Cardinal Mechanisms of Perception”) and a European Research Council (ERC) starting grant (ERC-2022-STG 101076057). This research was further supported by “The Adaptive Mind”, funded by the Excellence Program of the Hessian Ministry of Higher Education, Science, Research and Art. The authors declare that there are no competing interests.

## Supplementary Information

**Figure S1.**
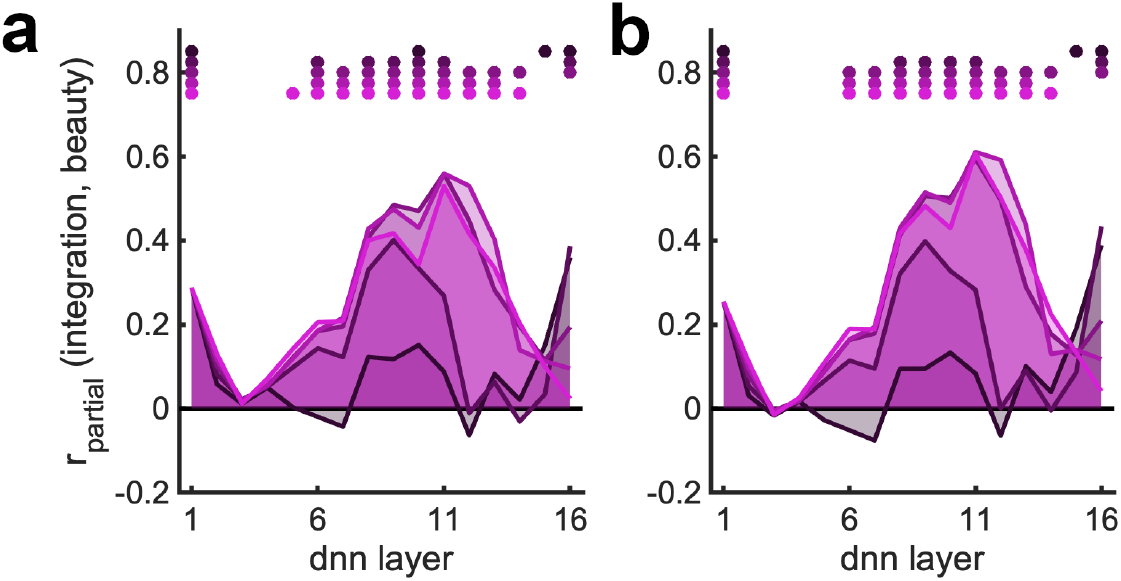
Behavioral ratings of complexity and order do not explain the correspondence between integration and perceived beauty. In Study 3, participants also rated the images for their complexity and order. **a)** Partialing out the complexity ratings did not account for the prediction of perceived beauty by our integration measure. **b)** Partialing out the order ratings did not account for the effect either. The same conventions apply as in Figure 2.

**Figure S2.**
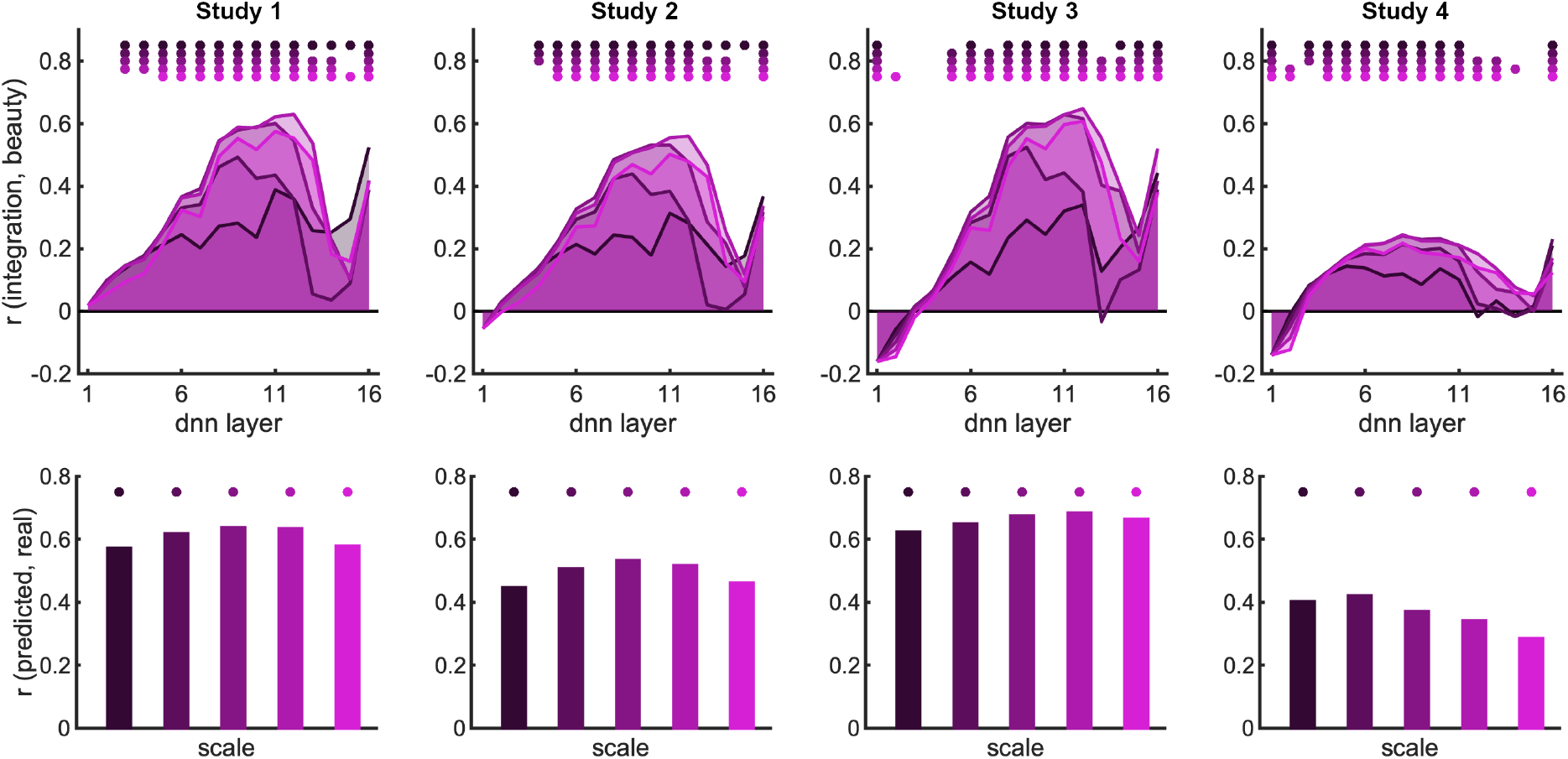
Integrative processing in an object categorization DNN also predicts perceived beauty. Similar predictions to the DNN trained on scene categorization were obtained with a DNN trained on object categorization (using the ImageNet image set). This suggests that integration occurs over features that are extracted both during object and scene categorization. It is, however, worth noting that the ImageNet dataset contains images of objects on scene backgrounds so that object and scene information cannot be fully orthogonalized using these networks. The same conventions apply as in Figure 2.

**Figure S3.**
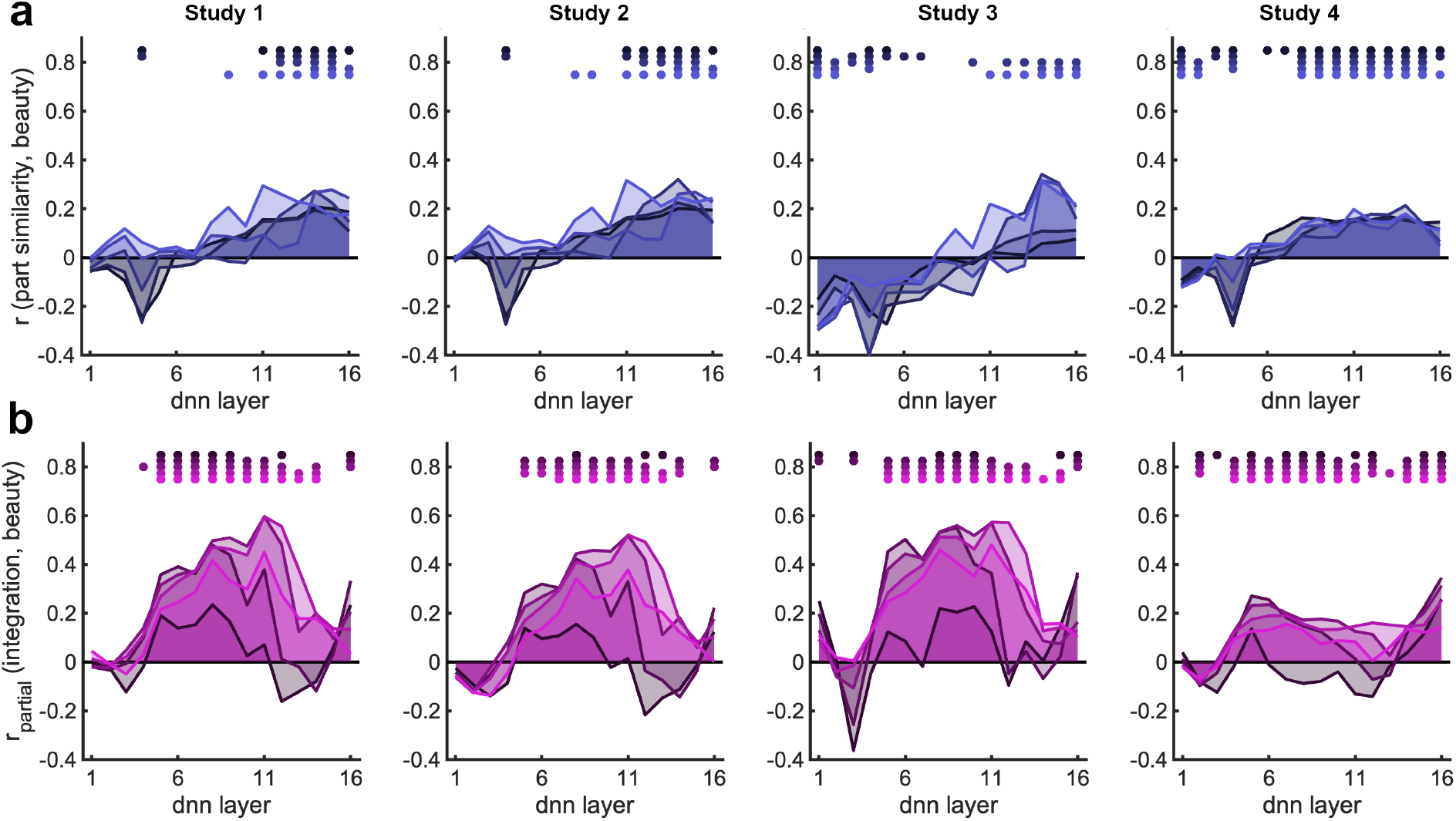
Part-based similarity also predicts perceived beauty but does not account for predictions based on integration. **a)** Like the integration measure, the part-based similarity measure (i.e., the similarity among image halves) also predicted beauty ratings across all studies. Interestingly, there was a difference between early in late network layers: the more similar the two halves are coded in the early layers, the lower their beauty rating, but the more similar they are coded in late layers, the higher their beauty rating. This suggests that diversity in low level features, but homogeneity in high-level features, makes images attractive. **b)** When partialing out the part-based similarity measure, however, our integration measure still significantly predicted perceived beauty. Same conventions apply as in Figure 2.

**Figure S4.**
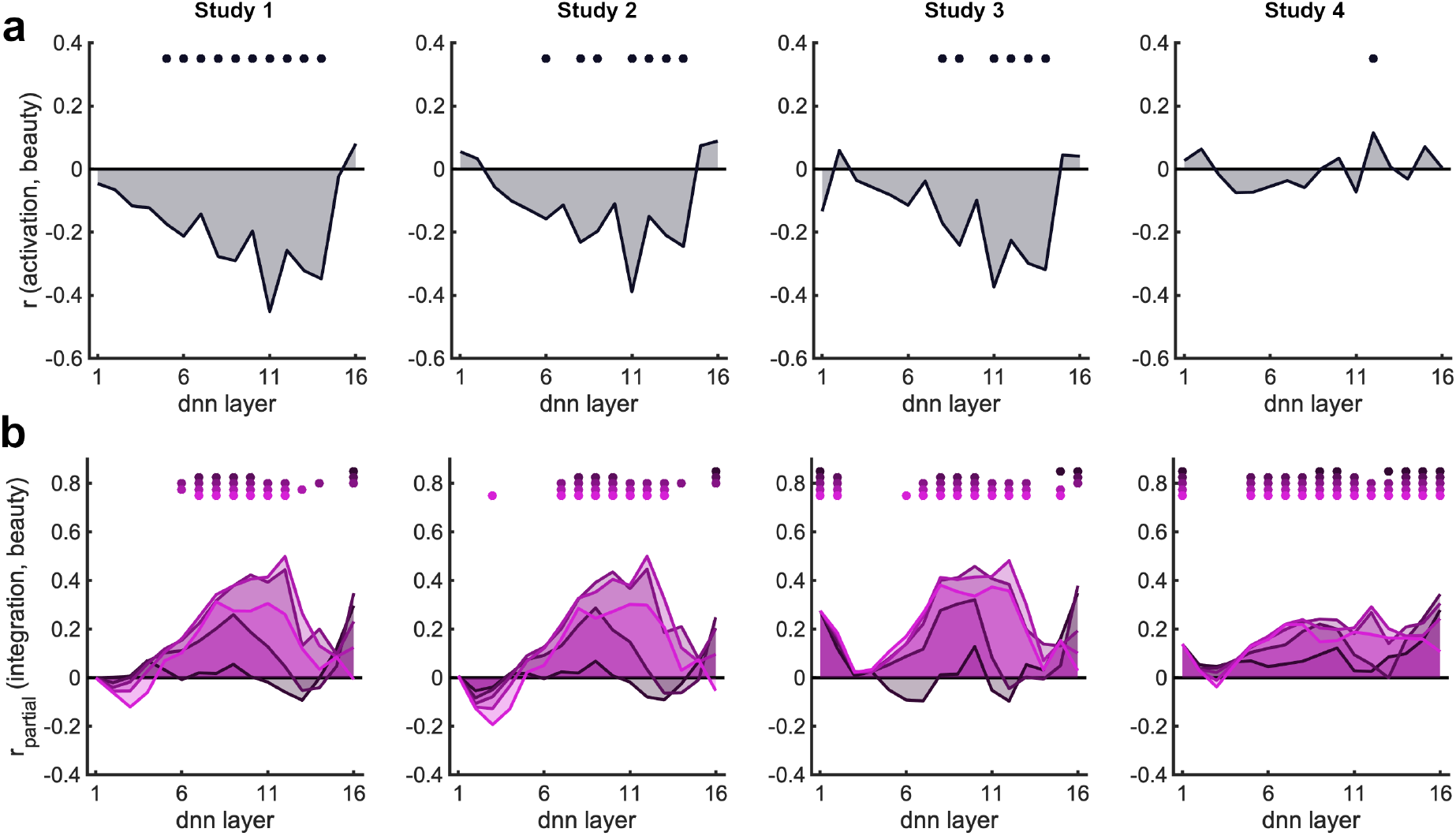
Activation strength (L2 norm) for the whole image also predicts perceived beauty but does not account for predictions based on integration. **a)** In Studies 1-3, images with lower activation strength in intermediate to late layers yielded higher beauty ratings. Such a relationship was not observed in Study 4. **b)** When partialing out the activation strength in response to the whole image, our integration measure still significantly predicted perceived beauty. Same conventions apply as in Figure 2.

